# Identification of representative trees in random forests based on a new tree-based distance measure

**DOI:** 10.1101/2022.05.15.492004

**Authors:** Björn-Hergen Laabs von Holt, Ana Westenberger, Inke R. König

## Abstract

In life sciences random forests are often used to train predictive models. However, gaining any explanatory insight into the mechanics leading to a specific outcome is rather complex, which impedes the implementation of random forests into clinical practice. By simplifying a complex ensemble of decision trees to a single most representative tree, it is assumed to be possible to observe common tree structures, the importance of specific features and variable interactions. Thus, representative trees could also help to understand interactions between genetic variants. Intuitively, representative trees are those with the minimal distance to all other trees, which requires a proper definition of the distance between two trees. Thus, we developed a new tree-based distance measure, which incorporates more of the underlying tree structure than other metrics. We compared our new method with the existing metrics in an extensive simulation study and applied it to predict the age at onset based on a set of genetic risk factors in a clinical data set. In our simulation study we were able to show the advantages of our weighted splitting variable approach. Our real data application revealed that representative trees are not only able to replicate the results from a recent genome-wide association study, but also can give additional explanations of the genetic mechanisms. Finally, we implemented all compared distance measures in R and made them publicly available in the R package timbR (https://github.com/imbs-hl/timbR).

## 1 Introduction

Predicting future outcomes for patients is a central aspect of modern medicine. Therefore, accurate and robust prognostic and predictive models are needed to predict, for instance, which individual treatment leads to the highest success probability. However, with the gaining importance of precision medicine and accompanying phenotype-rich data sets, the underlying mechanisms become too complex to handle with classical regression-based models. Thus, machine learning approaches become attractive.

One established method of machine learning are random forests which have become generally popular due to several advantages. First, they can be trained on a wide range of low- and high-dimensional data. Also, they can be applied in classification, regression as well as probability estimation settings (Breiman, 2001) and can even be used to model survival outcomes (Ishwaran *et al*., 2008; Hothorn *et al*., 2006). Finally, by aggregating single decision trees into a random forest, the prediction performance can usually be drastically improved. However, whereas single decision trees are easy to interpret, it is relatively difficult to understand and communicate the prediction by a random forest. Thus, similar to other complex machine learning approaches, random forests are often described to be predictive and explanatory black boxes. This often hinders their translation into clinical practice, given that it is more likely for a clinician to base his or her decision on a prediction model he or she understands. Hence, one of the key features of the usability of prediction models is that they are easy to interpret (Wyatt and Altman, 1995; Heinze *et al*., 2018).

To gain insight into a random forest model, a standard approach is to utilize variable importance measures (Breiman, 2001; Strobl *et al*., 2007; Nembrini *et al*., 2018). These evaluate how important each independent variable is for the performance of the model. However, this does not capture the complex tree structure of the model, including interactions between variables.

A completely different way to enhance the interpretability of a random forest is to identify a single tree, which best represents the forest. Thus, instead of theoretically having to inspect hundreds of trees in the entire forest, only one *representative tree* needs to be interpreted to understand the overall model. Banerjee *et al*. (2012) proposed to select those representative trees based on the highest similarity to all other trees in the forest. To measure the pairwise similarity of trees, they derived three distance metrics which cover different aspects of similarity of trees, namely, the similarity of predictions, the clustering in the terminal nodes, and the selection of splitting variables. However, these metrics are relatively simple and do not, for example, capture the underlying tree structure. Also, it should not only be of interest whether a variable was selected for splitting, but also at which point in the tree it was selected.

We therefore developed a new distance measure for decision trees to identify the most representative trees in random forests, based on the selected splitting variables but incorporating the level at which they were selected within the tree. We additionally incorporate the prioritization of variables that are selected multiple times in a tree, assuming that these are more important.

Section 2 of this paper begins with a short introduction to decision trees and random forests followed by a description of the metrics introduced by Banerjee *et al*. (2012). We then develop our new weighting splitting variable (WSV) metric and describe how to extract the most representative tree from the forest based on any tree distance. In Section 3, we investigate the behavior of the different tree-based distance measures by an extensive simulation study. We apply our new method to a real data set analyzing the genetic background of X-linked dystonia-parkinsonism (XDP) in Section 4. Therefore, we aim to predict the age at onset (AAO) of XDP patients by known genetic risk factors (Westenberger *et al*. (2019), Laabs *et al*. 2021). Finally, we discuss our results and give recommendations on the use of the different metrics in Section 5.

## 2 Methods

### Decision trees and random forests

Random forests (Breiman, 2001) are based on a combination of classification and regression trees (CART, Breiman *et al*., 1984) and Bagging (Breiman, 1996). They are a supervised learning algorithm meaning that the true outcome is known for all observations in the training data set. As a general setting, we assume a model to predict an outcome *y*, which may be categorical, ordinal or continuous, by a set of independent variables *x*_*p*_(*p* = 1, …, *P*).

To train a random forest model, a training data set *D* containing outcomes *y*_*n*_(*n* = 1, …, *N*) and independent variables *x*_*n,p*_ of *n* observations is used. From this, *num*.*trees* subsets *D*_*b*_, (*b* = 1, … *num*.*trees*) of *D* are generated by bootstrapping or subsampling. Based on every subset *D*_*b*_, a decision tree is trained by iteratively splitting *D*_*b*_ into binary subgroups. Unlike in the original CART algorithm, every split is based on just a subset of size *mtry* of the available independent variables, to increase the dissimilarity of the trees in a forest. Beginning with *D*_*b*_ acting as the root of the tree *T*_*b*_, the number of possible splits is given by the combination of splitting variables and splitting values. For each possible split *s* the decrease of impurity

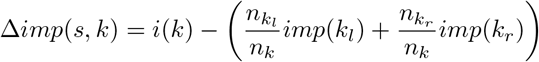

is calculated. Here, *imp*(*k*) is any impurity measure, and 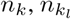 and 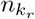 are the number of observations in node *k*, the left *k*_*l*_ and right *k*_*r*_ child node of *k*. For a categorical outcome variable *y* usually the Gini index is used as impurity measure, while the mean squared error (MSE) is used for continuous outcomes. Based on this, the optimal split *s*^*^ is the split which maximizes Δ*imp*(*s, k*). The training algorithm stops when either all leaves (terminal nodes) are pure or a pre-defined stopping criterion (e.g., minimal node size, maximal complexity, etc.) is fulfilled. The outcome of a new sample can be predicted by dropping the sample down the trees. Finally, the predictions of the *num*.*trees* trees are aggregated by majority vote for classification and by mean for regression trees, respectively.

### Distance Measures by Banerjee *et al*. (2012)

To select a limited subset of representative trees from the forest, Banerjee *et al*. (2012) derived three tree-based distance measures based on different aspects of the tree structure:

1. **Similarity of predictions (***d*_*pred*_): This compares the arbitrarily scaled predictions *ŷ*_*i,n*_ and *ŷ*_*j,n*_ of two decision trees *T*_*i*_ and *T*_*j*_ for observation *n* of an independent test data set, leading to the distance metric

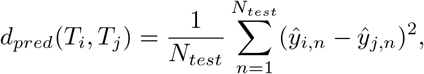

where *N*_*test*_ is the number of observations in the test data set. By comparing the predictions of trees this distance measure has a focus on selecting the tree with the most similar predictions to the complete ensemble, while it not necessarily gives the best explanation of the hidden tree mechanics.
2. **Similarity of clustering in terminal nodes (***d*_*clust*_): This evaluates whether two samples of an independent test data set end in the same leaf of *T*_*j*_ if they ended in the same leave of *T*_*i*_, leading to the metric

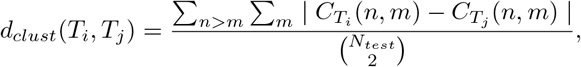

with

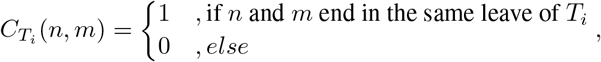

where *n* and *m* are observations of the test data set. In contrast to the first metric this one focus on how the trees separate the feature space, leading to a most representative tree that has the biggest overlap in the sparation of the feature space and thus a high explanatory power. But like the first metric an independent data set is necessary to estimate the distance making it highly dependent on the kind of data set available for calculations.
3. **Similarity of splitting variables (***d*_*split*_): This distance of two trees *T*_*i*_ and *T*_*j*_ is defined as

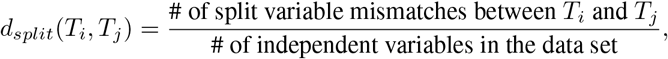

where one split variable mismatch occurs for an independent variable if one of the trees *T*_*i*_ and *T*_*j*_ uses this variable for splitting and the other does not. In contrast to the aforementioned methods the similarity of independent splitting variables can be estimated directly from each tree and thus does not need an independent data set making this method promising when no such data set is available. But since it only compares whether a variable is selected or not, its ability to cover the complex architecture of decision trees where a variable can be selected multiple times is limited.

In Figure 1 we built an example to highlight the differences between these methods. For this, we assumed a simple model with two dichotomous variables *x*_1_ and *x*_2_ explaining a dichotomous outcome *y*. Two non-identical trees *T*_1_ and *T*_2_ (upper half of Figure 1) were constructed and used to predict the class of four observations *o*_*n*_ = (*y*_*n*_, *x*_1*n*_, *x*_2*n*_), *o*_1_ = (−, 0, 0), *o*_2_ = (−, 1, 0), *o*_3_ = (+, 0, 1) and *o*_4_ = (−, 1, 1) (lower half of Figure 1). The pairwise tree distances for all three metrics as given above are shown. For *d*_*pred*_, the predictions are identical in both trees for all observations *o*_1_, …, *o*_4_, leading to a distance of 0. For *d*_*clust*_, we observe that the observation *o*_2_ ends in a terminal node together with *o*_4_ in *T*_1_ and together with *o*_1_ in *T*_2_. Thus, there are two differences in clustering, while we have 6 possible combinations according to the metric, leading to a distance of 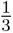. The metric *d*_*split*_ finally leads to a distance of 0, since both trees use the same split variable, but in a different way, which is ignored by *d*_*split*_.

**Figure 1:**
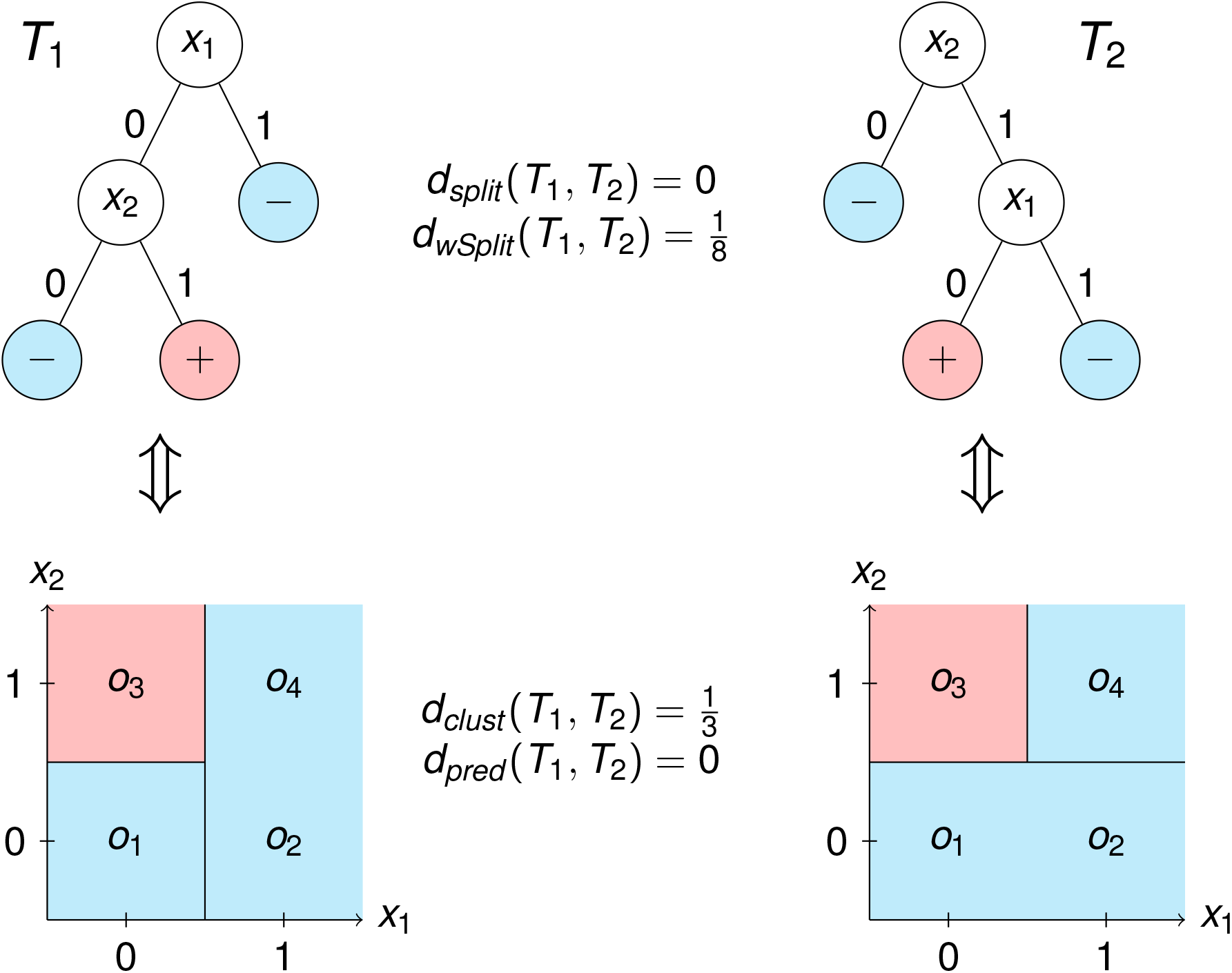
Comparison of the estimated pairwise tree distances based on four different metrics. Two trees *T*_1_ and *T*_2_ were constructed which use two dichotomous predictor variables *x*_1_ and *x*_2_ to predict a dichotomous outcome variable. Since both trees use exactly the same splitting variables, *d*_*split*_ is zero, although both trees use the two different variables in different ways. Therefore, *d*_*W SV*_ takes into account on which level of the tree a variable was used, resulting in a distance of 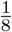. Based on a constructed test data set consisting of four observations *o*_*i*_ = (*y*_*i*_, *x*_1*i*_, *x*_2*i*_), *o*_1_ = (−, 0, 0), *o*_2_ = (−, 1, 0), *o*_3_ = (+, 0, 1) and *o*_4_ = (−, 1, 1), *d*_*pred*_ also leads to a distance of zero, since both trees give the same predictions on the test data set. Finally, the clustering by the two trees differ in whether *o*_2_ ends in a terminal node with *o*_1_ or *o*_4_, leading to a distance *d*_*clust*_ of 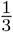

This example illustrates that the metrics *d*_*pred*_ and *d*_*split*_ proposed by Banerjee *et al*. (2012) can lead to a distance of 0 even for non-identical trees, because both do not exploit the tree structure in its entire complexity. The last metric *d*_*clust*_ uses the underlying tree structure for calculation but requires an independent test data set.

### Similarity of weighted splitting variable (WSV) (*d*_*W SV*_)

The WSV metric compares the structures of the trees in more detail without using an independent test data set. Specifically, we combine the information that a splitting variable *v* is used with the information about how much the tree relies on this splitting variable. The latter is assessed by a usage score

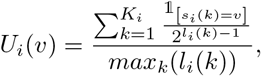

where *k* indicates the *K*_*i*_ nodes of *T*_*i*_. This leads to the distance measure

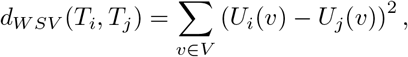

where *V* is the set of all independent variables. Thus, the WSV metric counts the number of times a specific splitting variable is used within a tree, weighted by the depth it occurs in. The weight becomes smaller the later in the tree the splitting variable is used, so that the first splitting variable receives the highest weight, assuming that the first split is the most important. The resulting vector of usage scores of all splitting variables is compared between two trees. Returning to the example in Figure 1, our new distance measure leads to usage scores 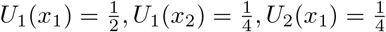 and 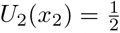, resulting in a distance of 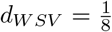. Therefore, this new metric is promising when no additional data set is available for estimating tree distances, since it covers more of the complex tree architecture of decision trees.

### Selection of the most representative tree

All of the tree-based distance metrics defined above measure the pairwise distances of trees based on different aspects of similarity. Given a specific distance metric, Banerjee *et al*. (2012) proposed to select the most representative tree from the random forest according to the distance score

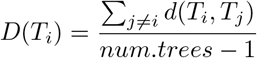

of every tree *T*_*i*_. From all trees, the tree with the lowest *D*(*T*_*i*_) is selected.

## 3 Simulation study

### Simulation design

Aim of our simulation study was to evaluate the properties of representative trees selected from a random forest based on different distance measures. For this, we defined the following data generating mechanism: We assumed a continuous outcome *y* which can be expressed by 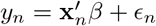, where *y*_*n*_ is the true outcome of observation *n*, and **x**_*n*_ = (*x*_*n,p*_)_=1,…,100_ ∼ ℬ (0.5) are independent variables with the effect sizes *β*_*p*_ and a normally distributed error term *ϵ*_*n*_ ∼ 𝒩 (0, 1). For each replicate of our simulations we simulated a training data set of 1000 observations (*y*_*n*_, *x*_*n*_) for training of random forests, a test data set of 100 observations for the calculation of distance metrics requiring independent test data and an additional validation data set of 1000 observations to compare tree and random forest predictions *ŷ*_*n*_ with the simulated *y*_*n*_.

We defined five different scenarios to be evaluated in our simulation study:

1. **Few large main effects:** In this most simple scenario only five independent variable have a *β*_*p*_ = 2, while all 95 other independent variables have an effect of zero.
2. **Many small main effects:** In contrast to the first scenario, here 50 independent variables have an effect *β*_*p*_ = 0.2 and the remaining 50 independent variable have no influence on the outcome *y*.
3. **Correlated variables:** While in the first two scenarios all independent variables were uncorrelated, we included correlated independent variables in the third scenario. Here, again we used five independent variables with *β*_*p*_ = 2 but this time, for each of the effect variables four additional variables were simulated which have a correlation of 0.3 to the effect variable, but no effect on the outcome themselves. Therefore, the complete data set consists of five variables with an effect, 20 correlated variables without main effects and 75 uncorrelated variables without an influence on the outcome.
4. **Interaction effects:** In this scenario again five independent variables have *β*_*p*_ = 2, but this time five additional pairs of variables have an interaction effect of the same size, but no main effect.
5. **Binary and continuous variables:** While in all other scenarios all independent variables were binary, we added five continuous variables sampled from a standard normal distribution to the data set alongside with five binary variables all having *β*_*p*_ = 2. Finally, the data set was filled with 90 binary variables without any influence on the outcome.

For each of the five scenarios we simulated 100 replicates. On each resulting training data set we trained four different random forests using ranger (Wright & Ziegler, 2017) with bootstrapping, 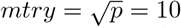 and varying minimal node sizes of ten, 50, 100 and 200. For every random forest, all four distance measures defined above were calculated, and the most representative trees were selected. Additionally, we trained single decision trees using rpart with the same minimal node sizes of ten, 50, 100, 200.

To assess the properties of the distance measures and resulting representative trees, we defined three different estimands with accompanying performance measures.

1. **Representation of the random forest prediction**: Here, we assess the ability of the most representative trees to represent the prediction of the entire ensemble. We selected for each metric the most representative tree with the lowest distance to all other trees, and evaluated the similarities of the predictions of these trees with those of the entire random forest by

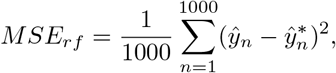

where *ŷ*_*n*_ is the prediction of the complete random forest on the validation data set, while 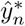 is the prediction of the selected most representative tree on the same data set.
2. **Generalizability of most representative trees**: Here, we evaluate the prediction performance of the most representative tree on a new independent data set. In the validation data the prediction accuracy of a most representative tree is investigated and compared with the accuracy based on the entire random forest. For this, we selected a most representative tree for each distance measure and compared its mean squared error with that of the entire random forest.
3. **Fraction of covered effect variables**: Ideally the selected most representative tree includes all effect variables. Therefore, we estimated how many of the simulated effect variables were at least once included in the most representative tree.

### Results on the representation of the random forest prediction

In Figure 2 the results for the representation of the random forest prediction is shown for all five scenarios. In all settings, the predictions of the most representative trees become more similar to those of the entire forest with increasing minimal node sizes. This is related to the fact that a larger minimal node size leads to smaller trees, which partition the prediction space more coarsely, so that a single tree can cover the information of the forest more easily. Additionally, the prediction based distance measure leads to the most similar predictions in all scenarios. This is also not surprising, since the distance measure uses the similarity of predictions defining the tree as most representative that is most similar to all other trees regarding their predictions. All other distance measures lead to rather similar results, but the WSV approach is the best of the remaining three metrics. In general most representative trees are more similar to the random forest than a decision tree that was trained on the same data set; but in a data set with many small main effects, the decision tree leads to remarkably better results than the most representative trees.

**Figure 2:**
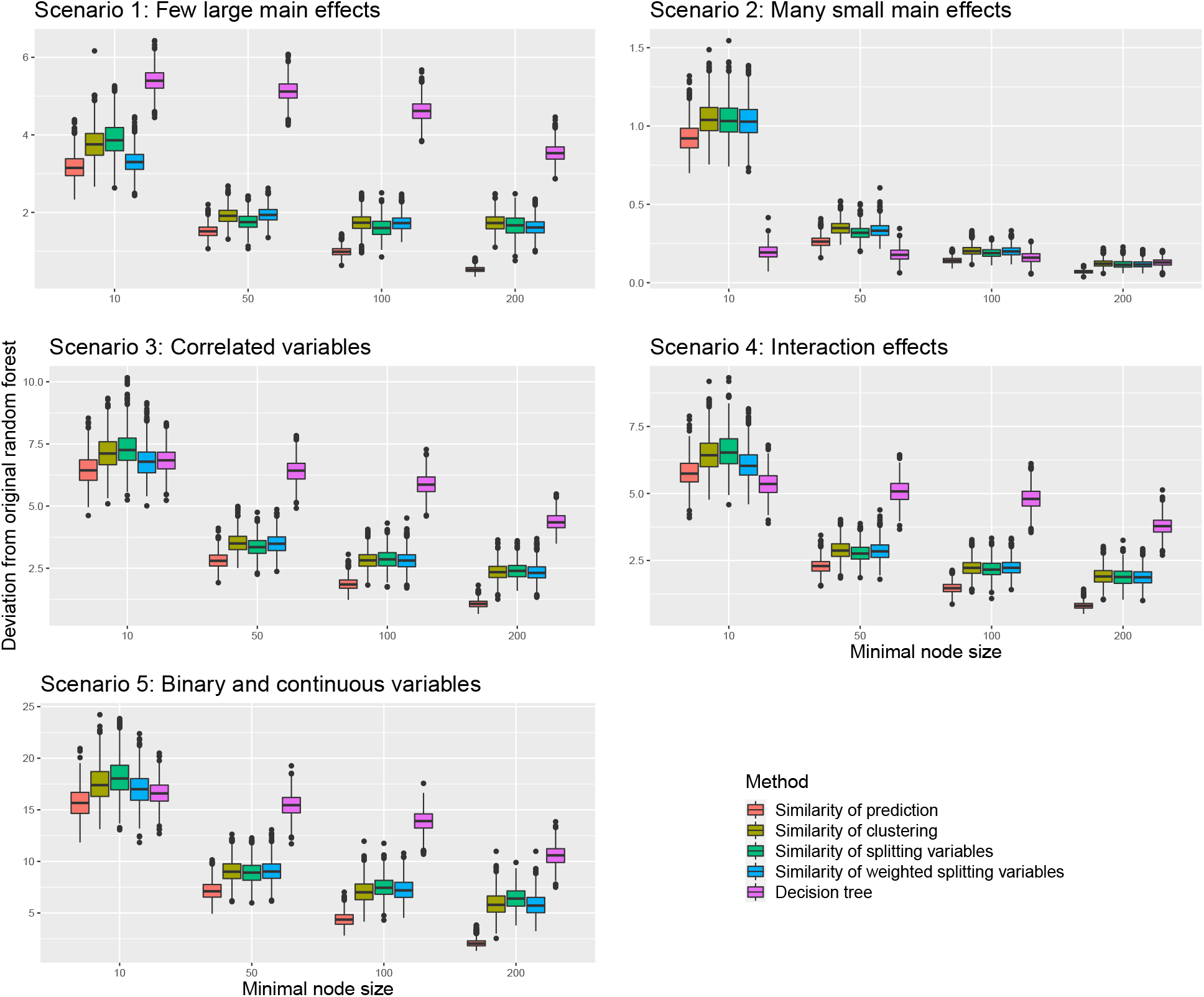
Representation of random forest predictions by single most representative trees selected based on four different distance measures, with respect to the minimal node size of the original random forest. Deviation of the representative tree prediction from the original random forest prediction was measured by the mean squared difference between the prediction of a most representative tree and the prediction of the complete random forest based on the same validation data set

### Results on the generalizability of most representative trees

The relative mean squared error of the most representative tree on a new validation data set is depicted in Figure 3. In scenario one, the WSV approach leads to the best MSE when the minimal node size is small and the trees are therefore more complex. For minimal node sizes above one hundred the distance measure using the clustering in the terminal nodes becomes slightly better. Of note, the prediction based distance measure is the second best approach for the very small minimal node size of ten, but is the worst for all other minimal node sizes. In contrast to this, the prediction based distance measure leads to the best results in scenario two, where many small effects influence the outcome, while all other metrics lead to similar results, that are slightly worse than the prediction based metric. When correlated independent variables or interaction effects are in the data set (scenario three and four), the WSV approach is again the best for random forests with a minimal node size of ten and the second best for higher minimal node sizes. In contrast to scenario one, this time the splitting variable approach is the best for higher minimal node sizes, but all but the prediction based approach are comparatively good. Finally, when a mixture of binary and continuous variables have an effect on the outcome, the prediction based approach leads to the best results when the minimal node size is ten followed by the WSV approach. In all other cases, the WSV distance is again only the second best, but the splitting variable distance yields the best results. Again, except for scenario two, most representative trees lead to better results than a single decision tree.

**Figure 3:**
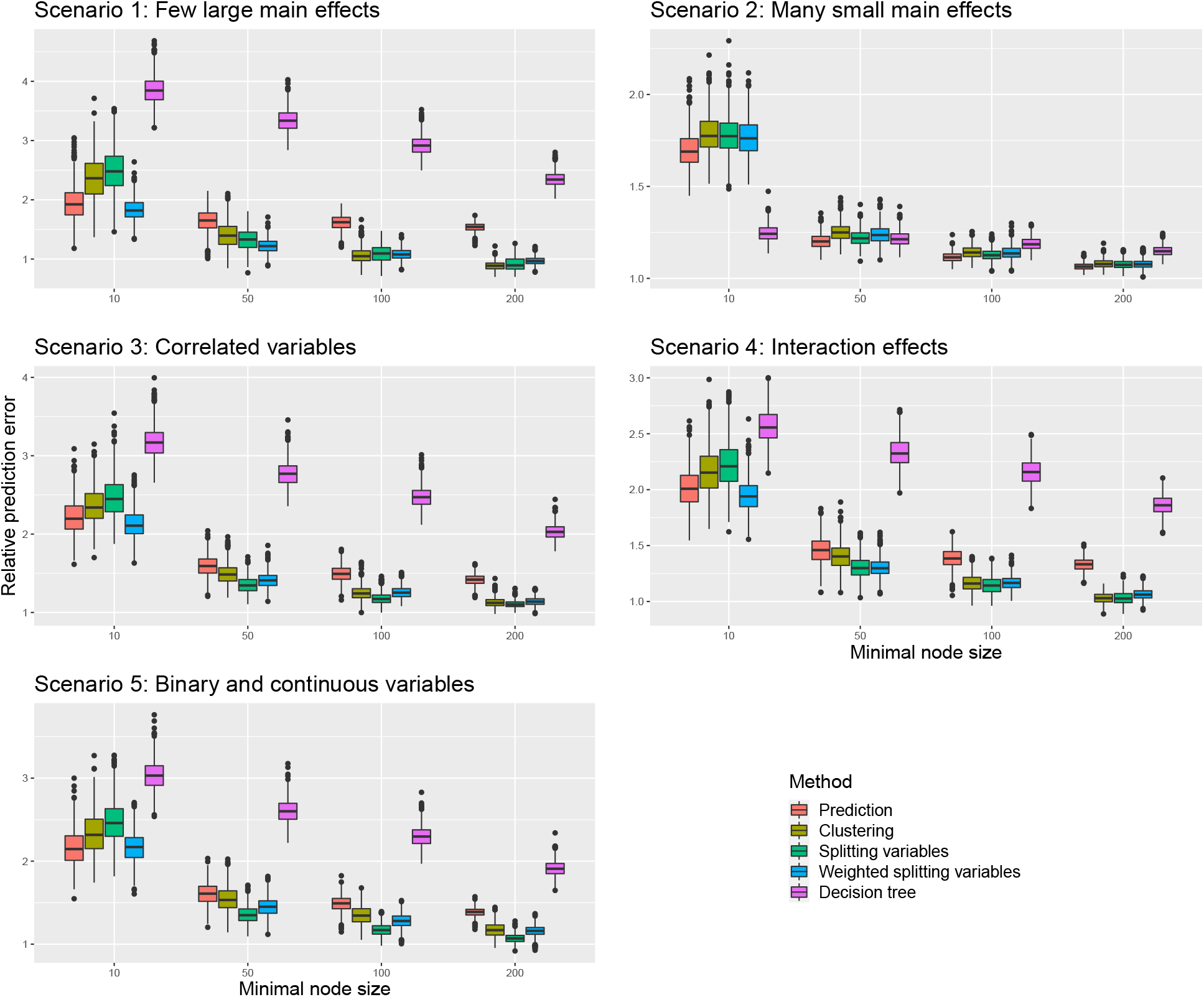
MSE of the prediction of the most representative tree selected by each method (color) on an independent test data set with respect to the minimal node size of the original random forest

### Computation time

While the computation time for each metric was constant in the different settings, the methods differ drastically. Specifically, the splitting variable distance is the fastest and only need 0.87 seconds to calculate, while the WSV metric is only slightly slower with 1.88 seconds. The prediction based metric is also comparatively fast with 2.93 seconds, while the clustering based approach is far slower with 488 seconds to calculate all pair-wise distances for 500 trees.

### Proportion of covered effect variables

Figure 4 depicts which mean proportion of the simulated effect variables were also included in the single generated trees. In scenario one, all trees cover all effect variables for a minimal node size of ten, but while the single decision tree still covers 90% of the effect variables for a minimal node size of 200, representative trees selected based on similarity of predictions drop below 20% for the same minimal node size. Overall it is observable that the similarity of prediction leads to representative trees that usually cover the least effect variables, which indicates that these trees may be good for prediction but not for explanation. Also, the single decision trees seem to have a lower coverage for many small main effects, interaction effects and the combination of continuous and dichotomous variables. Again, the similarity of weighted splitting variables leads to the trees with the second best coverage, but no other method performs constantly better.

**Figure 4:**
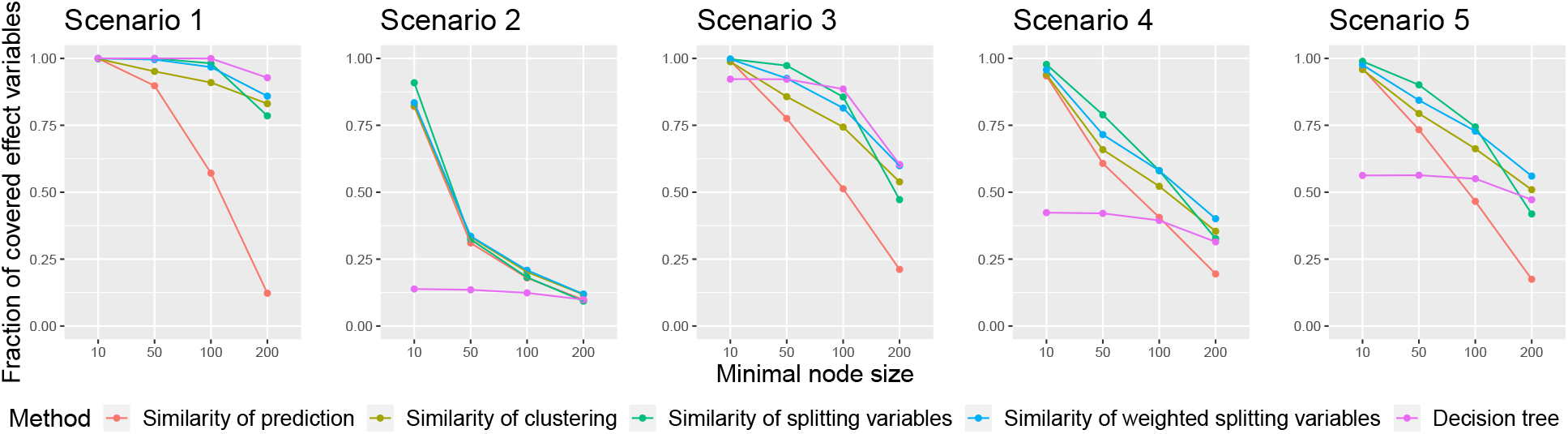
Proportion of simulated effect variables, that were covered by a single tree

## 4 Real data application

### Data set

To apply our newly developed distance measure to real data, we used data from 351 genetically confirmed XDP patients which was provided by the ProtectMove Research Unit (DFG: FOR2488). Participant enrollment and data analysis were approved by the Ethics Committees of the University of Lübeck, Germany, Massachusetts General Hospital, Boston, USA, Metropolitan Medical Center, Manila, Philippines, and Jose Reyes Medical Center, Manila, Philippines. XDP is a neurodegenerative movement disorder which combines symptoms of Dystonia and Parkinson’s disease and is limited to the Philippines (Lee *et al*. 2011). Whereas the genetic cause of XDP is well-understood (Aneichyk *et al*. 2018, Rakovic *et al*. 2018), the aim of current research is to better understand basis of the varying age at onset (AAO) in the affected patients. Two recent studies (Bragg *et al*. 2017, Westenberger *et al*. 2019) revealed that 50% of the variability in AAO can be explained by the repeat number of a hexanucleotide repeat (RN) within the ∼2,6-kb *SINE-VNTR-Alu (SVA)* retrotransposon insertion in intron 32 of *TAF1* on the X chromosome. In our study population the AAO ranges between 21 and 67 years (mean = 41.7 years, standard deviation (sd) = 8.4 years) and RN ranges between 30 and 55 units (mean = 41.6 units, sd = 4.1 units). For all patients the AAO was determined in a standardized interview with movement disorder specialists. For each observation the data set includes not only AAO and RN, but also a set of 82 additional single nucleotide polymorphisms (SNPs). Therefore, we selected three SNPs which were recently identified to modify AAO in XDP (Laabs *et al*. 2021) located in *MSH3* and *PMS2*. Additionally, we selected the SNPs with the lowest p-values in the genome-wide association study by Laabs *et al*. (2021) within nine different genes in the *MSH3/PMS2* pathway. Since XDP patients show symptoms similar to dystonia (DYT) and Parkinson’s disease (PD), we also included the SNPs with the lowest p-values in 12 PD and 12 DYT genes as well as 36 PD-related SNPs (Nalls *et al*. 2019) and 8 DYT-related SNPs (Mok *et al*. 2014, Lohmann *et al*. 2014). Finally, two SNPs associated with Huntington’s disease (HD) were included (Moss *et al*. 2017, Chao *et al*. 2018), since Laabs *et al*. 2021 revealed a possible relationship between HD and XDP. Details to collection and processing of the gentic data can be found in Laabs *et al*. 2021. In total this leads to a data set of 351 samples and 83 independent variables.

### Real data results

We aimed to extract a single most representative tree for the given problem of modeling AAO in XDP patients and to examine its tree structure. Therefore, we trained a single random forest on the complete available data set with a number of trees of 500, *mtry* = *P* (so that on each node the best variable can be selected), bootstrapping and a minimal node size of 20% of the training data set. From the resulting random forest we selected the most representative tree (Figure 5). Here, the first four splits were all made at the SVA RN, separating the observations into different groups with increasing RN. This is not surprising, since Westenberger *et al*. (2019) described a negative linear influence of RN on the AAO. In further concordance with Westenberger *et al*. (2019), the predicted AAO for the groups with smaller RN were also higher. Interestingly, for the group with the lowest RN (≤ 36.5 units), no further genetic variant was selected for splitting. Another surprising fact was that the three variants reported by Laabs *et al*. (2021) never occurred together in one branch of the tree. The variant rs33003 in *MSH3* was only used for splitting in patients with lower RN (*>* 36.5 but ≤ 39.5 units), while the other variant in *MSH3* (rs245013) was selected for splitting in patients with higher RN (*>* 39.5 but ≤ 43.5 units). Finally, the variant in *PMS2* (rs62456190) was of importance for patients with RN between 43.5 and 50.5 units. For the patients with the highest RN (*>* 50.5 units), again no additional genetic information was used besides the SVA RN. After the split on rs245013, the tree uses variants not identified by Laabs *et al*. (2021) for the first and only time. Here we can see that in patients who were homozygous for the alternative allele of rs11060180, a variant in the *CCDC62* gene which was previously associated with Parkinson’s disease, was associated woth a high decrease in AAO, making it a potential risk factor. The fact that all known SNPs associated with the AAO of XDP were selected in different branches of the tree indicates that they tag modifiers that do not interact with each other, but act individually. Additionally, it is likely, that each of the tagged modifiers interacts with the RN.

**Figure 5:**
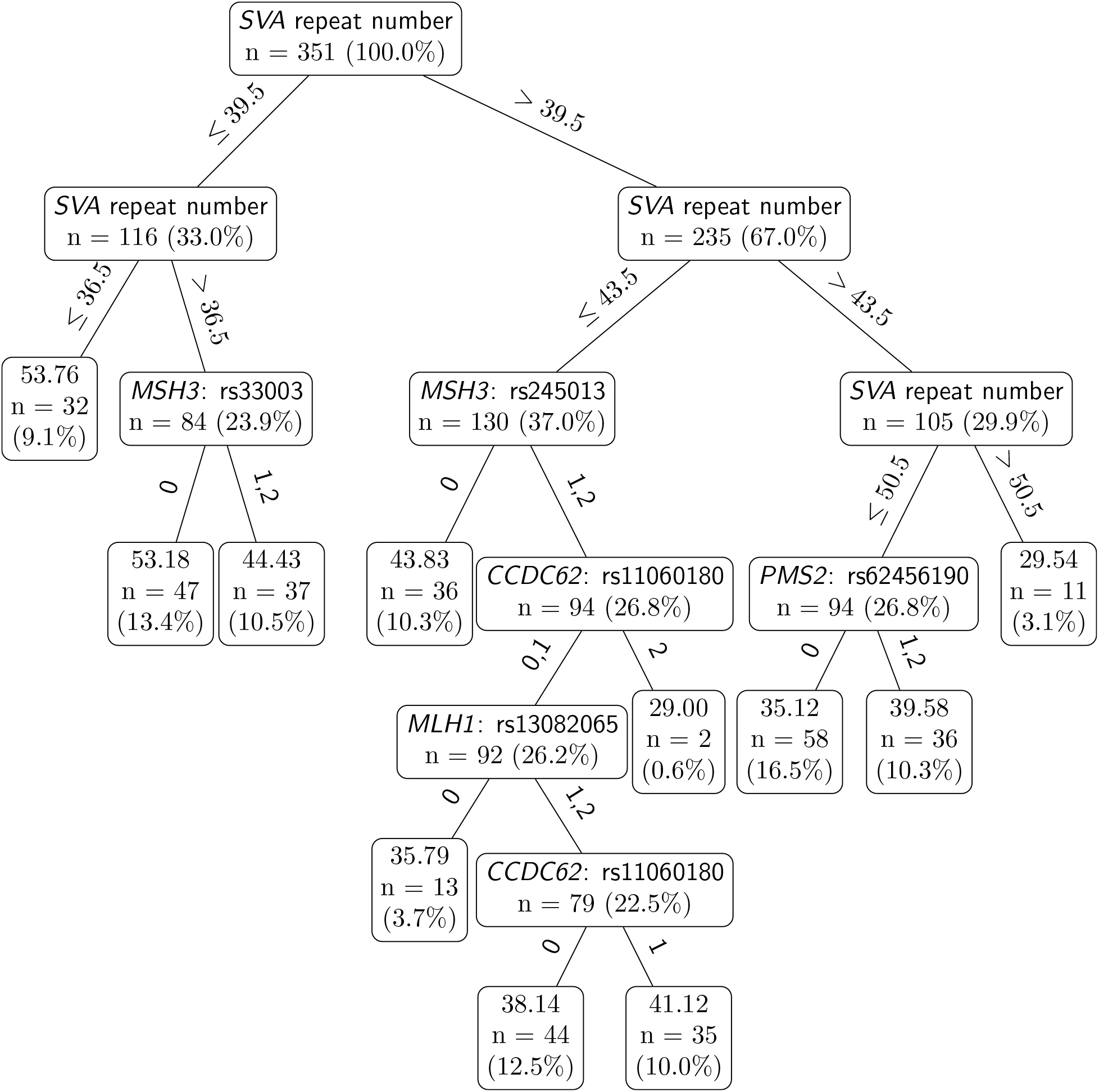
Most representative tree selected by our newly developed distance metric

## 5 Discussion

Our work aimed to develop a new tree-based distance measure capable of using the underlying tree architecture to estimate the similarity of decision trees. This is necessary to select most representative trees from random forests, which have a considerable potential in the visualization and interpretation of random forests and may facilitate the incorporation of machine learning into clinical practice. Therefore, we constructed a usage score which estimates how a given tree uses a specific variable by increasing the score for each occurrence of the variable in the tree weighted by the level of the tree it occurs in. Thus, the earlier a variable is selected for splitting, the higher the score. Our distance metric then compares the usage score for all variables between two trees and summarizes the squared difference of them.

To investigate the validity of the WSV metric in comparison with other distance measures, we designed an extensive simulation study in which we considered five different data scenarios. In these, we compared the prediction of the most representative tree with the complete random forest prediction. Here, the prediction based metric by Banerjee *et al*. (2012) that requires an additional test data for estimating the distance performed best, but the WSV metric performed second best in most scenarios. Additionally, we compared the prediction of the most representative tree with the known true outcome of the observations in an additional validation data set. In this case, the prediction based distance metric leads to the best results when the minimal node size is very small and the trees are therefore more complex. In all other cases the distance measures using the splitting variables or the clustering in terminal nodes lead to the best results. Of note, the WSV approach was again the second best approach in most of the cases.

Although the most representative trees selected by the WSV metric are only the second-best in nearly all scenarios, they showed the most robust results, while each other metric sometimes lead to the best and sometimes to the worst results. Another advantage of the WSV approach is that it does not rely on an additional data set to estimate distances, like other metrics, making it valuable for situations without additional available data. Of note, the addtional test data sets generated in our simulation study used the exact same generating mechanism as the original training data set. In practice it can be expected that an additional data set underlies a slightly different generating mechanism leading to an expected decrease of the explanatory power of most representative trees that rely on such data sets. Finally, WSV is comparably fast and leads reliably to the trees which include the most effect variables.

In an application on real data predicting the AAO of XDP patients, we were able to replicate the results by Westenberger *et al*. 2018 and Laabs *et al*. 2021. As a major advantage, our approach takes the position of a variable within the tree into account and is, therefore, better able to mirror the tree architecture. However, it still does not incorporate the sequence or interaction of variables. Thus, further studies that may show how the WSV distance metric may be improved in a time-efficient way are warranted. We expect, that the improvement in the calculation of pair-wise tree distances are further steps to improve the interpretability and therefore the practical use of random forests.

## Acknowledgments

This work was supported by the Deutsche Forschungsgemeinschaft (DFG; FOR 2488 to IRK and AW).

## References

Aneichyk T. Hendriks W.T., Yadav R., Shin, D., Gao, D., Vaine, C.A., Collins, R.L., Domingo, A., Currall, B., Stortchevoi, A., Multhaupt-Buell, T., Penney, E.B., Cruz, L., Dhakal, J., Brand, H., Hanscom, C., Antolik, C., Dy, M., Ragavendra, A., Underwood, J., Cantsilieris, S., Munson, K.M., Eichler, E.E., Acun∼a, P., Go, C., Jamora, R.D.G., Rosales, R.L., Church, D.M., Williams, S.R., Garcia, S., Klein, C., Müller, U., Wilhelmsen, K.C., Timmers, H.T.M., Sapir, Y., Wainger, B.J., Henderson, D., Ito, N., Weisenfeld, N., Jaffe, D., Sharma, N., Braekefield, X.O., Ozelius, L.J., Bragg, D.C. and Talkowski, M.E. (2018). Dissecting the Causal Mechanism of X-Linked Dystonia-Parkinsonism by Integrating Genome and Transcriptome Assembly. Cell 172(5), 897–909 https://doi.org/10.1016/j.cell.2018.02.011.

Banerjee, M., Ding, Y. and Noone, A.-M. (2012). Identifying representative trees from ensembles. Statistics in Medicine 31(15), 1601–1616 https://doi.org/10.1002/sim.4492.

Bragg, D.C., Mangkalaphiban, K., Vaine, C.A., Kulkarni, N.J., Shin, D., Yadav, R., Dhakal, J., Ton, M.L., Cheng, A., Russo, C.T., Ang, M., Acu∼na, P., Go, C., Franceour, T.N., Multhaupt-Buell, T., Ito, N., Müller, U., Hendriks, W.T., Breakfield, X.O., Sharma, N. and Ozelius, L.J. (2017). Disease onset in X-linked dystonia-parkinsonism correlates with expansion of a hexameric repeat within an SVA retrotransposon in TAF1. Proceedings of the National Academy of Sciences of the United States of America 114(51), E11020–E11028 https://doi.org/10.1073/pnas.1712526114.

Breiman L. (1996). Bagging predictors. Machine Learning 24(2), 123–140 https://doi.org/10.1007/BF00058655.

Breiman L. (2001). Random forests. Machine Learning 45(1), 5–32 https://doi.org/10.1023/A:1010933404324.

Breiman, L., Friedman, J., Stone, C.J. and Olshen, R.A. (1984). Classification and regression trees. Chapman & Hall/CRC, Boca Raton.

Chao, M.J., Kim, K.-H., Shin, J.W., Lucente, D., Wheeler, V.C., Li, H., Roach, J.C., Hood, L., Wexler, N.S., Jardim, L.B., Holmans, P., Jones, L., Orth, M., Kwak, S., MacDonald, M.E., Gusella, J.F. and Lee, J.-M. (2018). Population-specific genetic modification of Huntington’s disease in Venezuela. PLOS Genetics 14(5), https://doi.org/10.1371/journal.pgen.1007274.

Heinze, G., Wallisch, C. and Dunkler, D. (2018). Variable selection - A review and recommendations for the practicing statistician. Biomtrical Journal 60(3), 431–449 https://doi.org/10.1002/bimj.201700067.

Hothorn, T., Hornik, K. and Zeileis, A. (2006). Unbiased recursive partitioning: A conditional inference framework. Journal of Computational and Graphical Statistics 15(3), 651–674 https://doi.org/10.1198/106186006X133933.

Ishwaran, H., Kogalur, U.B., Blackstone, E.H. and Lauer, M.S. (2008). Random survival forests. The Annals of Applied Statistics 2(3), 841–860 https://doi.org/10.1214/08-AOAS169.

Laabs, B.-H., Klein, C., Pozojevic, J., Domingo, A., Brüggemann, N., Grütz, K., Rosales, R.L., Jamora, R.D., Saranza, G., Diesta, C.C.E., Schaake, S., Dulovic-Mahlow, M., Quismundo, J., Otto, P., Acuna, P., Go, C., Sharma, N., Multhaupt-Buell, T., Müeller, U., Hanssen, H., Kilpert, F., Rolfs, A., Bauer, P., Dobricic, V., Lohmann, K., Ozelius, L.J., Kaiser, F.J., König, I.R. and Westenberger, A. (2021). Identifying novel genetic modifiers of age-associated penetrance in X-linked dystonia-parkinsonism. Nature Communications 12, 3216.

Lee l.V., Rivera, C., Teleg, R.A., Dantes, M.B., Pasco, P.M.D., Jamora, R.D.G., Arancillo, J., Villareal-Jordan, R.F., Rosales, R.L., Demaisip, C., Maranon, E., Peralta, O., Borres, R., Tolentino, C., Monding, M.J. and Sarcia, S. (2011). The unique phenomenology of sex-linked dystonia parkinsonism (XDP, DYT3, “Lubag”). International Journal of Neuroscience 121(1), 3–11 https://doi.org/10.3109/00207454.2010.526728.

Li, J., Mallay, J.D., Andrew, A.S., Karagas, M.R. and Moore, J.H. (2016). Detecting gene-gene interactions using a permutation-based random forest method. BioData Mining 9(14), 1–17 https://doi.org/10.1186/s13040-016-0093-5.

Lohmann K., Schmidt, A., Schillert, A., Winkler, S., Albanese, A., Baas, F., Bentivoglio, A.R., Borngräber, F., Brüggemann, N., Defazio, G., Del Sorbo, F., Deuschl, G., Edwards, M.J., Gasser, T., Gómez-Garre, P., Graf, J., Groen, J.L., Grünewald, A., Hagenah, J., Hemmelmann, C., Jabusch, H.C., Kaji, R., Kasten, M., Kawakami, H., Kostic, V.S., Ligouri, M., Mir, P., Münchau, A., Ricchiuti, F., Schreiber, S., Siegesmund, K., Svetel, M., Tijssen, M.A., Valente, E.M., Westenberger, A., Zeuner, K.E., Zittel, K.E., Altenmüller, E., Ziegler, A. and Klein, C. (2014). Genome-wide association study in musician’s dystonia: a risk variant at the arylsulfatase G locus? Movement Disorders 29(7), 921–927 https://doi.org/10.1002/mds.25791.

Mok, K.Y., Schneider, S.A., Trabzuni, D., Stamelou, M., Edwards, M., Kasperaviciute, D., Pickering-Brown, S., Silverdale, M., Hardy, J. and Bhatia, K.P. (2014). Genomewide association study in cervical dystonia demonstrates possible association with sodium leak channel. Movement Disorders 29(2), 245–251 https://doi.org/10.1002/mds.25732.

Moss, D.J.H., Langbehn, D., Leavitt, B.R., Roos, R., Durr, A., Mead, S., TRACK-HD investigators, REGISTRY investigators, Holmans, P., Jones, L. and Tabrizi, S.J. (2017). Identification of genetic variants associated with Huntington’s disease progression: a genome wide association study. Lancet Neurology 16(9), 701–711 https://doi.org/10.1016/S1474-4422(17)30161-8.

Nalls, M.A., Blauwendraar, C., Vallerga, C.L., Heilbron, K., Bandres-Ciga, S., Chang, D., Tan, M., Kia, D.A., Noyce, A.J., Xue, A., Bras, J., Young, E., von Coelln, R., Simón-Sánchez, J., Schulte, C., Sharma, M., Krohn, L., Pihlstrom, L., Siitonen, A., Iwaki, H., Leonard, H., Faghri, F., Gibbs, J.R., Hernandez, D.G., Scholz, S.W., Botia, J.A., Martinez, M., Corvol, J.-C., Lesage, S., Jankovic, J., Shulman, L.M., The 23andMe Research Team, System Genomics of Parkinson’s Disease Consortium, Sutherland, M., Tienari, P., Majamaa, K., Toft, M., Brice, A., Yang, J., Gon-Or, Z., Gasser, T., Heutink, P., Shulman, J.M., Wood, N., Hinds, D.A., Hardy, J., Morris, H.R., Gratten, J., Vischer, P.M., Graham, R.R. and Singleton, A.B. (2018). Parkinson’s disease genetics: identifying novel risk loci, providing causal insights and improving estimates of heritable risk. bioRxiv doi: https://doi.org/10.1101/388165

Nembrini, S., König, I.R. and Wright, M.N. (2018). The revival of the Gini importance?. Bioinformatics 34(21), 3711–3718 https://doi.org/10.1093/bioinformatics/bty373.

Rakovic, A., Domingo, A., Grütz, K., Kulikovskaja, L., Capetian, P., Cowley, S.A., Lenz, I., Brüggemann, N., Rosales, R., Jamora, D., Rolfs, A., Seibler, P., Westenberger, A., König, I., and Klein, C. (2018). Genome editing in induced pluripotent stem cells rescues TAF1 levels in X-linked dystonia-parkinsonism. Movement Disorders 33(7), 1108–1118 https://doi.org/10.1002/mds.27441.

Strobl, C., Boulesteix, A.-L., Zeileis, A. and Hothorn, T. (2007). Bias in random forest variable importance measures: Illustrations, sources and a solution. BMC Bioinformatics 8(25), 1–21 https://doi.org/10.1186/1471-2105-8-25.

Westenberger, A., Reyes, C.J., Saranza, G., Dobricic, V., Hanssen, H., Domingo, A., Laabs, B.-H., Schaake, S., Pozojevic, J., Rakovic, A., Grütz, K., Begemann, K., Walter, U., Dressler, D., Bauer, P., Rolfs, A., Münchau, A., Kaiser, F.J., Ozelius, L.J., Jamora, R.D., Rosales, R.L., Diesta, C.C.E., Lohmann, K., König, I.R., Brüggemann, N. and Klein, C. (2019). A hexanucleotide repeat modifies expressivity of X-linked dystonia parkinsonism. Annals of Neurology 85, 812–822 https://doi.org/10.1002/ana.25488.

Wright, M.N. and Ziegler, A. (2017). ranger: A fast implementation of random forests for high dimensional data in C++ and R. Journal of Statistical Software 77,1–17. https://doi.org/10.18637/jss.v077.i01.

Wyatt, J.C., and Altman D.G. (1995). Prognostic models: clinically useful or quickly forgotten? The BMJ 311, 1539–1541 https://doi.org/10.1136/bmj.311.7019.1539.

